# Skeletal muscle reprogramming by breast cancer regardless of treatment history or tumor molecular subtype

**DOI:** 10.1101/810952

**Authors:** Hannah E. Wilson, David A. Stanton, Cortney Montgomery, Aniello M. Infante, Matthew Taylor, Hannah Hazard-Jenkins, Elena N. Pugacheva, Emidio E. Pistilli

## Abstract

Increased susceptibility to fatigue is a negative predictor of survival commonly experienced by women with breast cancer. Here, we sought to identify molecular changes induced in human skeletal muscle by BC regardless of treatment history or tumor molecular subtype using RNA-sequencing and proteomic analyses. Mitochondrial dysfunction was apparent across all molecular subtypes, with the greatest degree of transcriptomic changes occurring in women with HER2/neu-overexpressing tumors, though muscle from patients of all subtypes exhibited similar pathway-level dysregulation. Interestingly, we found no relationship between anti-cancer treatments and muscle gene expression, suggesting that fatigue is a product of BC *per se* rather than clinical history. *In vitro* and *in vivo* experimentation confirmed the ability of BC cells to alter mitochondrial function and ATP content in muscle. These data suggest that interventions supporting muscle in the presence of BC-induced mitochondrial dysfunction may alleviate fatigue and improve the lives of women with BC.

## INTRODUCTION

Muscle dysfunction in individuals with cancer is commonly thought to be a consequence of muscle atrophy, which is a major component of the paraneoplastic syndrome known as cancer cachexia ^1, 2^. While studies in men with cancer support the claim that muscle functional capacity is dependent on muscle size, women with cancer report a significant degree of muscle dysfunction despite typically remaining weight-stable ^3–5^. Muscle dysfunction in breast cancer (BC) commonly presents as a persistent, severe fatigue that frequently contributes to dose-reduction or treatment cessation and is an independent predictor of survival in a variety of cancer types, including BC ^6–11^. Thus, it is probable that improving muscle fatigue will improve both quality of life and survival in BC.

At present, there are several purported contributory factors for cancer-related fatigue, including immunological responses to tumor growth; side-effects of cancer therapies; depression and/or emotional distress; anemia; hormonal, nutritional, and metabolic disturbances; and inadequate physical activity ^8, 12^. Pharmacological treatments for cancer-related fatigue exhibit limited and inconsistent success, in part because determining the mechanisms contributing to fatigue in a given patient can be quite challenging, particularly in patients with early-stage disease and those not receiving anti-cancer treatments ^13^. Identifying mechanisms of BC-related fatigue that are intrinsic to skeletal muscle, generalizable across BC subtypes, and independent of treatment status could significantly aid in the development of appropriate therapies, which would be applicable to a large number of patients experiencing BC-associated fatigue.

Our laboratory has recently reported that skeletal muscle of women with BC exhibits a distinct gene expression signature that is not dependent on molecular subtype ^14^. Furthermore, we have identified signaling via the metabolic regulators of the peroxisome-proliferator activated receptor (PPAR) family as potential key mediators of fatigue in women with BC and female mice bearing BC patient-derived orthotopic xenografts (PDOXs) ^15^. Our previous analyses did not include women with primary tumors that overexpressed HER2/neu in the absence of estrogen receptor (ER) and progesterone receptor (PR) expression. In the current study, we have expanded our analyses into all molecular subtypes, including both transcriptomic and proteomic analyses of muscle biopsies from patients with HER2/neu-overexpressing tumors, and significantly increased our sample size to create, to our knowledge, the largest study of transcriptomic and proteomic changes in muscle of women with BC. We tested the hypothesis that BC induces a common molecular response in skeletal muscle that is independent of the molecular subtype of the tumor and the patient’s treatment history.

## RESULTS

### Patient characteristics

A total of 51 BC patients representing 4 breast tumor subtypes and 20 non-cancer controls provided *pectoralis major muscle* biopsies and/or detailed clinical information for use in the present study. There were no differences in mean body mass index (BMI) between non-cancer controls and BC patients; average BMI in the control group was categorized as Class I Obesity and in the BC patients was categorized as Overweight. Additionally, there were no significant differences in the percent (%) change in BMI, body fat %, or lean body mass between controls and BC patients. There were no differences in BMI between any of the 4 breast tumor subtypes (Table 1).

**Table 1:**
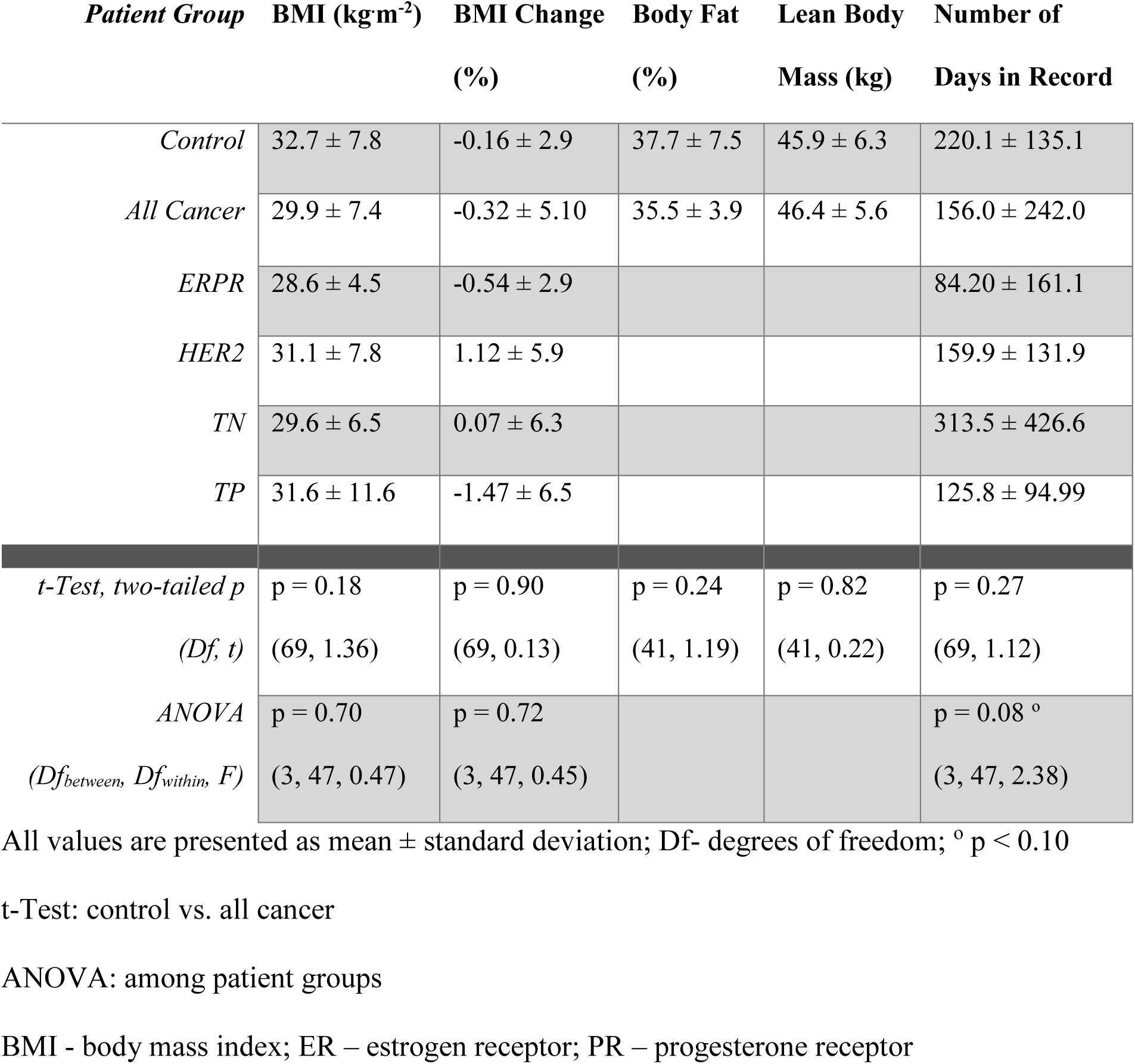
Patient Characteristics.

### Skeletal muscle gene expression profiles

Skeletal muscle biopsies from patients diagnosed with BC (n=33) and non-cancer controls (n=10) were used for RNA-seq, and BC patients were classified based on molecular subtype of their primary tumor, as follows: luminal (ERPR)— positive for estrogen receptor (ER) and progesterone receptor (PR) without overexpression of HER2/neu; HER2 —overexpression of HER2/neu in the absence of ER and PR expression; triple negative (TN)—absence of ER, PR, and HER2/neu expression; and triple positive (TP)— presence of ER and PR expression, and overexpression of HER2/neu (ERPR n=10, HER2 n=5, TN n=9, TP n=9). Unsupervised clustering analysis indicated a significant degree of clustering based on molecular subtype, particularly with regard to patients with tumors overexpressing HER2/neu in the absence of ER or PR (Figure 1A). Multidimensional scaling (MDS) analysis in 3 dimensions revealed that the gene expression profiles of skeletal muscle from patients with ERPR, TP and TN tumors were similar, while the profile from skeletal muscles from patients with HER2/neu-overexpressing tumors was significantly different (Figure 1B). Overall fit of the MDS model was dramatically improved by the use of BC subtype as a covariate (Figure 1B, adjusted R^2^=0.44, p=0.0001) rather than a model including only binary disease status (Figure 1C, adjusted R^2^=0.20, p=0.008), which was already a significant improvement over the null model (Figure 1D).

**Figure 1:**
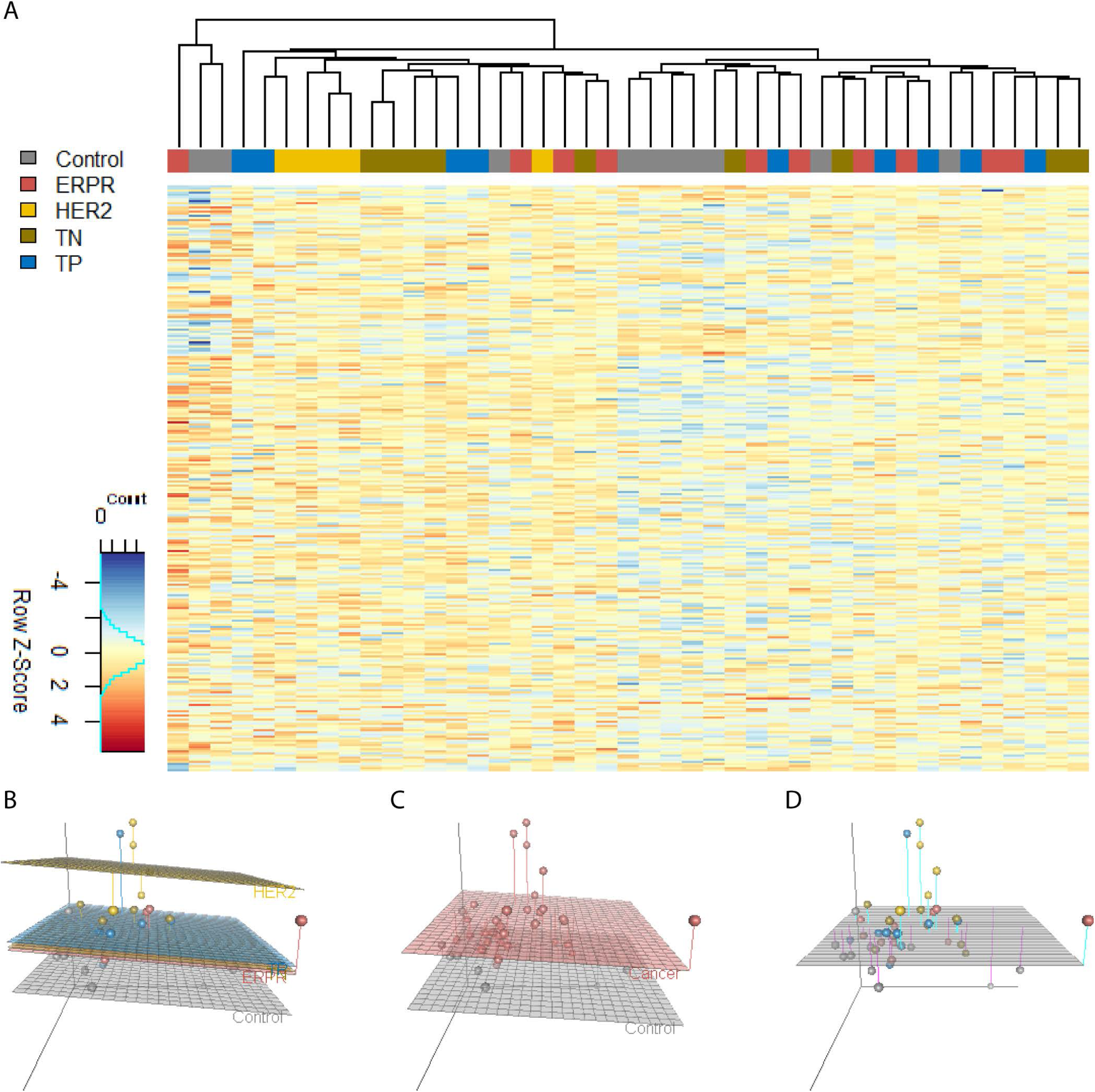
Skeletal Muscle Gene Expression Profiles. Unsupervised hierarchical clustering on normalized, log-transformed RNA-sequencing gene expression data of the 10,000 most variable genes across all patients (**A**). MDS dissimilarity matrix of overall gene expression data represented in 3 dimensions, with each dot representing an individual patient. Regression planes are color-coded based on molecular subtype (**B**), binary disease status (**C**), or the null model (**D**).

To address the question of whether this obvious difference in overall muscular gene expression between groups was inherent to differences in the primary tumor type or the myriad of clinical characteristics that could potentially differ between groups, we assessed the relationship between clinical characteristics and skeletal muscle gene expression in the context of a multivariate linear regression model, using the 3 dimensions of the MDS dissimilarity matrix as response variables. Among the various treatment types, body composition, serum albumin, and changes in body mass, the only assessed variable to yield statistical significance at α=0.05 when used as a single independent variable was patient group (i.e. Control, ERPR, HER2, TN, TP), with serum albumin nearing statistical significance (Table 2). Notably, chemotherapy, radiotherapy, and hormonal treatments did not correlate with overall gene expression patterns, nor did the patient’s trend of weight change over time. Using forward selection, a final model including patient group and serum albumin was identified as the best-fitting model for predicting muscular gene expression from clinical data (Table 3).

**Table 2:** Possible clinical predictors of muscular gene expression.

| *Variable* | **Variable Type**<br>*(Levels or Range)* | **Pillai's**<br>**Trace** | **Num**<br>**Df** | **Den**<br>**Df** | **p** |
| --- | --- | --- | --- | --- | --- |
| *Group* | Factor<br>*(Control, ERPR, HER2, TN, TP)* | 0.51 | 12 | 111 | 0.04 * |
| *Serum Albumin* | Numeric | 0.20 | 3 | 29 | 0.08 ° |
| *N Staging* | Factor *(N0, N1, N1a, N2, N3)* | 0.52 | 12 | 84 | 0.15 |
| *# of Chemotherapy TxS* | Numeric *(0-2)* | 0.10 | 3 | 38 | 0.25 |
| *# of Radiation TxS* | Numeric *(0-1)* | 0.10 | 3 | 38 | 0.26 |
| *T Staging* | Factor *(T1, T1a, T2, T2A, T3, T4)* | 0.51 | 15 | 81 | 0.37 |
| *# of Hormonal TxS* | Numeric *(0-1)* | 0.07 | 3 | 38 | 0.43 |
| *LBM %* | Numeric | 0.12 | 3 | 19 | 0.45 |
| *BMI* | Numeric | 0.05 | 3 | 38 | 0.56 |
| *Ave. Daily BMI Change* | Numeric | 0.03 | 3 | 37 | 0.78 |
| *M Staging* | Factor *(M0, MX)* | 0.01 | 3 | 29 | 0.95 |

Num- numerator; Den- denominator; Df- degrees of freedom; LBM %- Lean body mass as a percentage of total body mass; Ave- Average; T- tumor; N- lymph node; M- metastatic; TxS- courses of treatments; \* $p < 0.05$ , ° $p < 0.10$

**Table 3:** Final clinical predictors of muscular gene expression.

| *Model Formula* | **F-Statistics for Model Fit,**<br>**By Dimension** | **Pillai's Trace**<br>**(p-value)** | **Wilk's Lambda**<br>**(p-value)** |
| --- | --- | --- | --- |
| *(D1, D2, D3) ~ Patient Group* | D1: 0.3<br>D2: 6.9<br>D3: 0.4 | 0.506<br>(0.044) * | 0.522<br>(0.020) * |
| *(D1, D2, D3) ~ Patient Group*<br>*+ Serum Albumin* | D1: 0.7<br>D2: 3.5<br>D3: 2.1 | 0.744<br>(0.052) ° | 0.401<br>(0.049) * |

D1-D3- individual dimensions of RNA-sequencing data as 3-dimensional dissimilarity matrix;
\* $p < 0.05$ , ° $p < 0.10$

Because serum albumin appeared to provide predictive value for skeletal muscle gene expression data, and because serum albumin is commonly used by oncologists to monitor patients’ nutritional status, the relationship between serum albumin and changes in BMI over time were assessed in the group of patients that provided biopsies. There was no correlation observed between serum albumin at the date of biopsy collection and the individual’s rate of weight change over time **(Figure2A)**. Similar results were obtained in a retrospective chart review of 3,001 patients with BC. While there was a statistically significant correlation between a patient’s first record of serum albumin and the patient’s rate of weight change in this large sample (Figure 2B), the effect size may well be clinically insignificant (R = 0.094). The average daily weight change was negligible in women with normal serum albumin as well as those with low serum albumin (< 3.4 g **·** dL^-1^), with both groups having means within one standard deviation of 0 (Figure 2C). For a patient to be considered cachectic by traditional standards, an average daily weight loss of at least 0.027% would be required to lose 5% of their weight in 6 months ^1, 2^. In our large cohort, a logistic regression analysis was conducted using a threshold of 3.4 g **·** dL^-1^ serum albumin to predict whether a patient would exhibit this rate of weight change. In this analysis, omnibus model fit was significant at ɑ < 0.05, though effect size as determined by Nagelkerke’s pseudo-R^2^ was very small and indicates a very weak predictive value (R^2^ = 0.08). Additionally, a threshold of 3.4 g **·** dL^-1^ serum albumin was only 40% sensitive to identifying this level of weight change and only yielded a positive predictive value of 17.3%. In other words, 60% of BC patients exhibiting a rate of weight loss consistent with cachexia had normal serum albumin, while BC patients with serum albumin < 3.4 g **·** dL^-1^ only had a 17.3% chance of exhibiting a rate of weight change consistent with cachexia.

**Figure 2:**
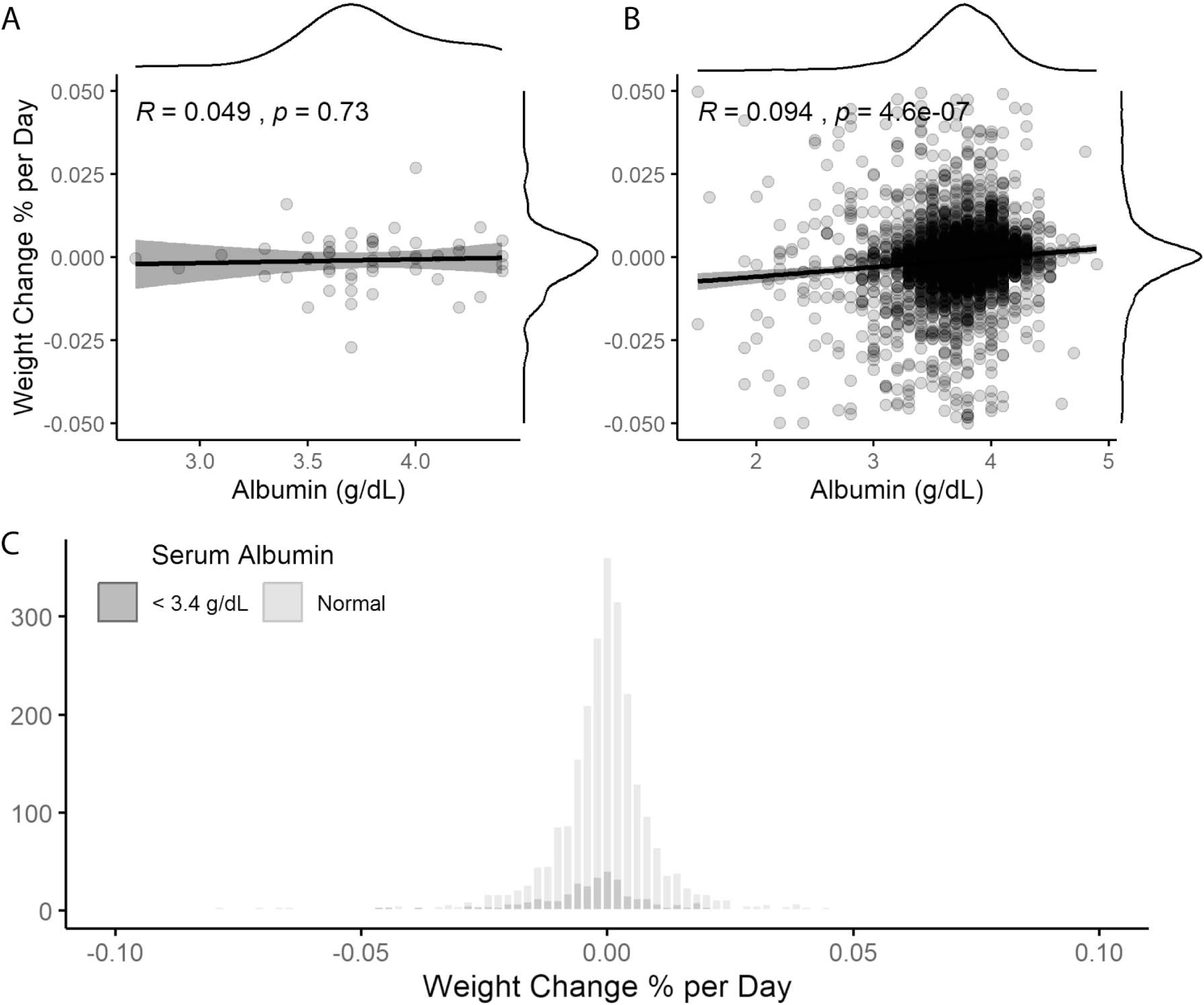
Serum Albumin and Weight Loss. Linear regression analysis of trends in weight change predicted by serum albumin in patients with BC who provided muscle biopsies and at least one serum albumin measurement. Grey dots represent individual patients; grey shading represents 95% confidence interval of the black regression line, with marginal kernel density plots provided for both variables (**A**, n=51). Linear regression analysis of trends in weight change predicted by serum albumin in a retrospective chart review of patients with BC. Grey dots represent individual patients; grey shading represents 95% confidence interval of the black regression line, with marginal kernel density plots provided for both variables (**B**, n=3,001). Histogram representing individualized rate of weight change in a retrospective chart review of patients with BC, with patients grouped by serum albumin level (**C**, n_low_=442, n_normal_=2,557).

### Differential gene expression analysis by subtype

Differentially expressed genes (DEGs) within skeletal muscle were first identified by comparing BC patients by subtype to control. Considerable overlap of DEGs was observed between the 4 breast tumor subtypes. Of the 3,468 genes identified as differentially expressed in at least one subtype, only 7 (0.2%) were unique to ERPR patients, 173 (5.0%) to TN patients, and 80 (2.3%) to TP patients. However, 2,410 genes (70%) were unique to HER2 patients, and only 8 (0.23%) were differentially expressed in all patient groups (Figures 3A and 3B). These observations were quantified and reveal that there was an approximately 2-fold fewer-than-expected number of unique DEGs in muscle of patients with ERPR, TP and TN tumors if the DEGs were independent of subtype. Specifically, one would expect 14, 186, and 508 unique DEGs in each subtype, respectively, whereas we actually observed 7, 80, and 173 DEGs in these groups. Further, HER2 patients’ muscle exhibited 2-fold fewer-than-expected DEGs shared with any combination of two other subtypes (Observed = 5 + 2 + 84 = 91; Expected total = 203; overall *χ*^2^ on 6 degrees of freedom = 1224, p = 3 x 10^-261^). This indicates that the ERPR, TP, and TN groups share a greater number of DEGs in skeletal muscle than one would expect if the DEGs were independent of subtype and in contrast, muscle from HER2 patients does not exhibit the same similarity to the other subtypes in terms of shared DEGs (Figure 3C). Collectively, these data demonstrate that transcriptional responses in skeletal muscle of patients with ERPR, TP and TN tumors are highly similar, in support of previous data from our laboratory ^14, 15^. Furthermore, the transcriptional responses in muscles from patients with HER2/neu-overexpressing tumors partially overlap with the other subtypes, but exhibit a significant contrast to the other 3 subtypes, suggesting that this tumor type is associated with a unique transcriptional adaptation within skeletal muscle.

**Figure 3:**
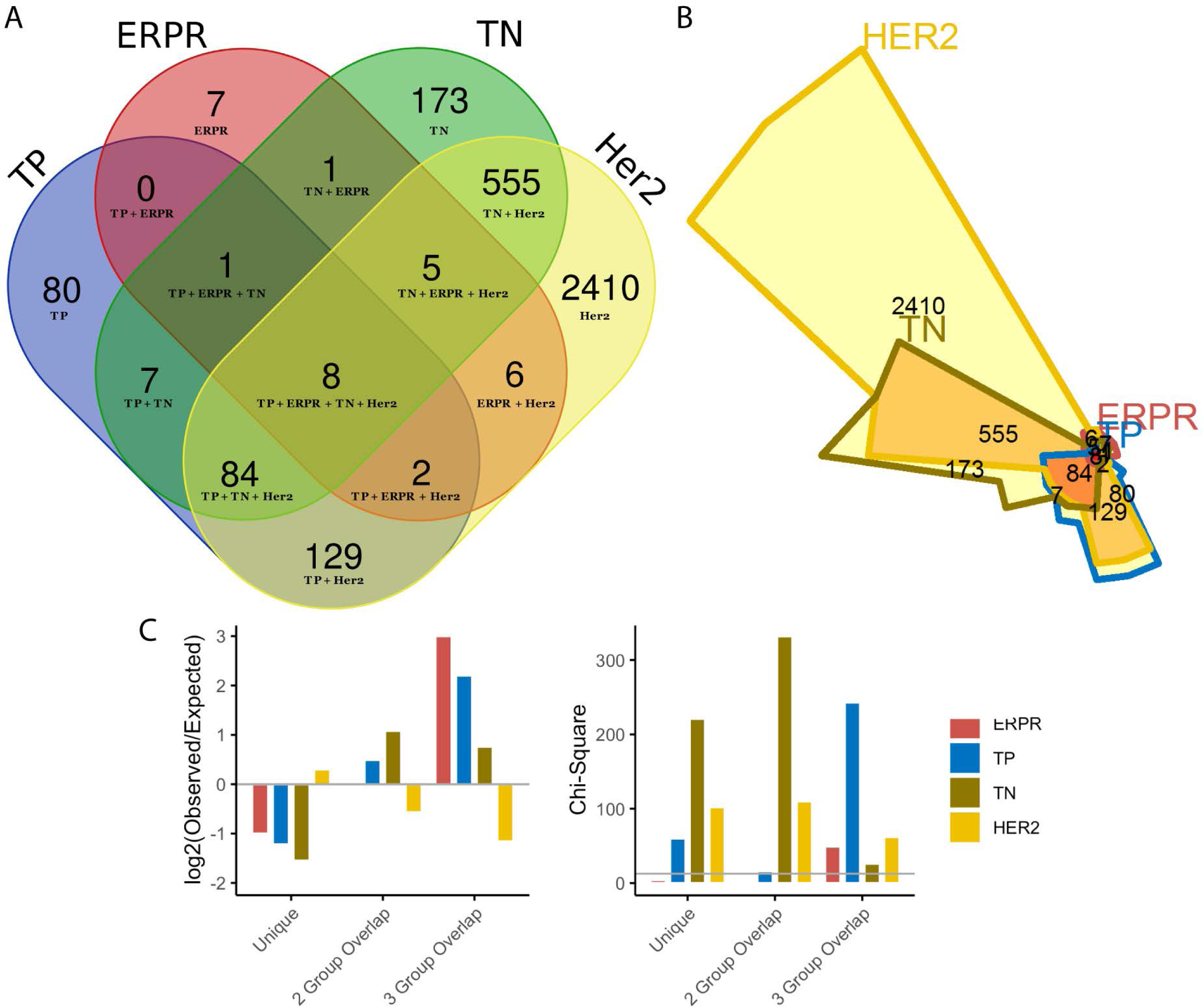
Differential Gene Expression Analysis by Subtype. Venn diagram representing DEGs in muscle in each molecular subtype (**A**) and corresponding area-representative Chow-Ruskey diagram (**B**). Chi-square analysis of the number of DEGs uniquely differentially-expressed in each subtype, the number of DEGs involved in 2- and 3-way overlaps among subtypes, showing the log-fold change between observed and expected numbers in each category The Chi-square critical value of 12.592 (6 degrees of freedom) is denoted on the Chi-Square y-axis by a gray horizontal line. A Chi-square value larger than 12.592 is statistically significant at α = 0.05. (**C**).

### DEG pathway analysis by subtype

Qiagen’s Ingenuity Pathway Analysis (IPA) software was used to infer pathway-level dysregulation based on trends in transcriptomic changes. Identified pathways were shared between all breast tumor subtypes to a greater extent than individual genes. Of the 368 pathways identified by IPA as dysregulated in any subtype, 43 (11.7%) pathways were identified as significantly dysregulated in all 4 breast tumor subtypes relative to control. 23 (6.3%) pathways were uniquely dysregulated in muscle from patients with TP tumors, 25 (6.8%) in patients with ERPR tumors, 29 (7.9%) unique to TN patients, and 62 (16.8 %) uniquely dysregulated in patients with HER2/neu-overexpressing tumors (Figures 4A, 4B, 4C). Chi-square analysis to test whether the number of pathways shared between subtypes differed between groups was non-significant (overall *χ*^2^ on 6 degrees of freedom = 8.10, p = 0.23), indicating that the four BC subtypes share a common core of dysregulated pathways. When the 43 overlapping pathways were ranked by how similarly all subtypes appeared in terms of magnitude and directionality of dysregulation, LXR/RXR signaling stood out in its strong, consistent inhibition across all BC subtypes (Figure 4D), and the related TR/RXR pathway was also identified as consistently significantly dysregulated. These pathways are closely related to and often integrated with the PPAR/RXR signaling pathway, which our laboratory has previously identified as a likely upstream regulator of muscle fatigue in BC patients ^14, 15^. Additional dysregulated pathways were related to inflammation and immune system processes.

**Figure 4:**
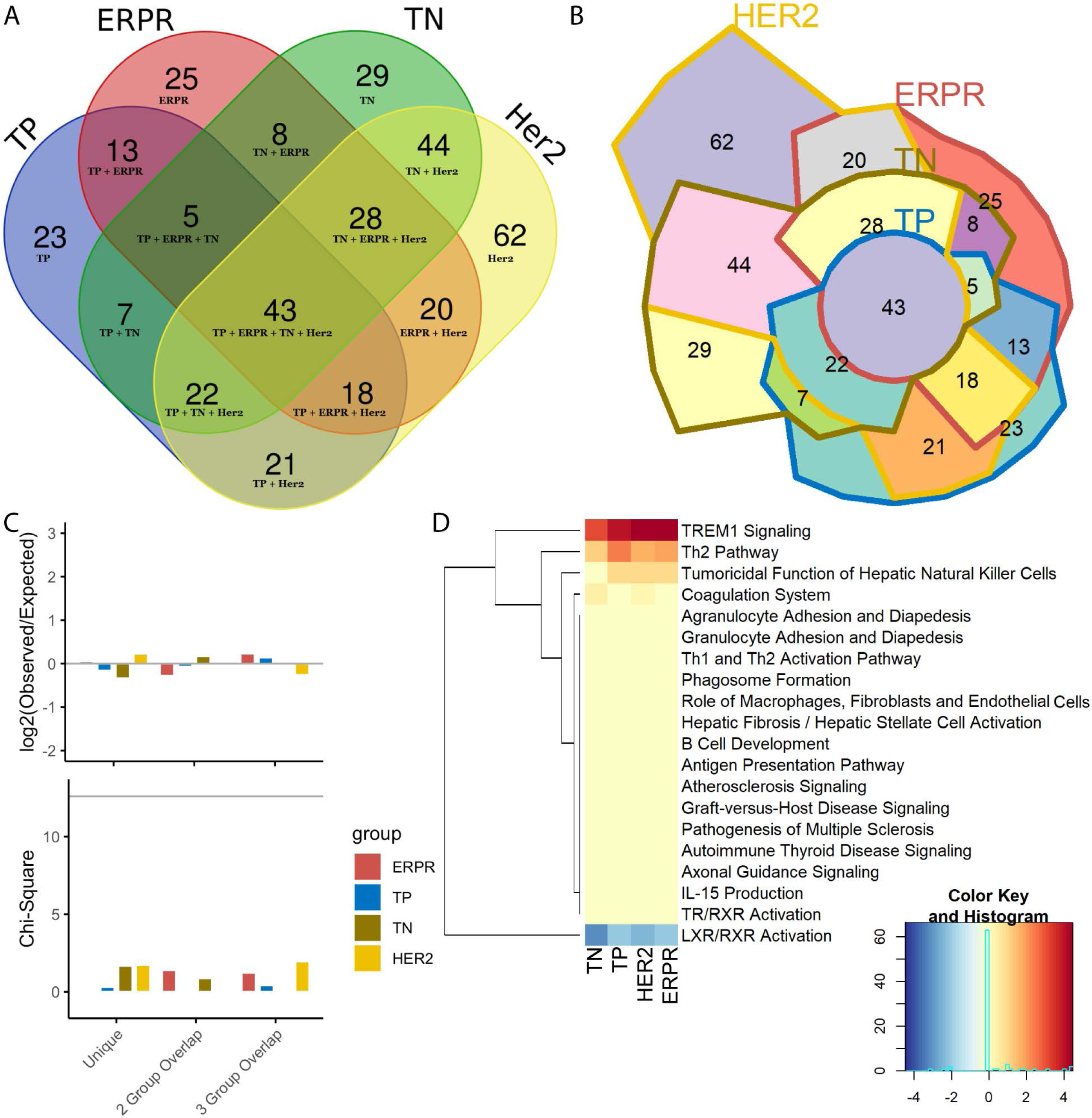
Dysregulated Pathway Analysis by Subtype. Venn diagram representing significantly dysregulated IPA pathways in muscle in each molecular subtype (**A**) and corresponding area-representative Chow-Ruskey diagram (**B**). Chi-square analysis of the number of pathways uniquely dysregualted in each subtype and those involved in 2- and 3-way overlaps between subtypes, showing the log-fold change between observed and expected numbers in each category. The Chi-square critical value of 12.592 (6 degrees of freedom) is denoted on the Chi-Square y-axis by a gray horizontal line. A Chi-square value larger than 12.592 is statistically significant at α = 0.05. (**C**). Heatmap of activity scores for the twenty most consistently dysregulated pathways across all subtypes (i.e. pathways with lowest variance in activity scores), with the four columns of the heatmap representing the four molecular subtypes. Lower activity scores in blue indicate pathway inhibition and higher activity scores in red indicate pathway activation (**D**).

### HER2 patient proteomics

Because the HER2 patient group exhibited the greatest degree of transcriptomic dysregulation, muscle biopsies from these patients (n=5) were selected for proteomic analysis and compared to control surgical patients (n=5). Scaled, log-transformed expression data from proteomic analysis were correlated with scaled, log-transformed RNA-sequencing expression data in a gene-wise fashion. 8/8 individuals with matched RNA-seq and proteomic analyses were found to have moderately strong correlation (Pearson’s R range 0.48 – 0.52, **Supplemental Figure 1**). A total of 1,555 unique proteins were detected across all samples, 1,259 of those at a high confidence level, with most proteins being detected in all 10 samples (Figure 5A). Differential expression analysis detected only a small number of significantly differentially expressed proteins (FDR < 0.05, Figure 5B), though there are obvious physiological implications in the small set. The six downregulated differentially expressed proteins (DEPs) were all identified as mitochondrial components (Figure 5C), representing a statistically significant enrichment of mitochondrial components in this small set of DEPs (Figure 5D, p = 3.2 x 10^-6^). Expanding our analysis to both significant DEPs and insignificant trends in protein expression, a strong signal for aberrant mitochondrial function was once again detected. Nearly every protein involved in the mitochondrial electron transport chain was quantified as having a lower level of protein expression in the HER2 patients relative to controls (Figure 5E**, Supplementary Table 1),** including proteins encoded by both nuclear DNA and mitochondrial DNA, and mitochondrial function and oxidative phosphorylation were identified as being the two most significantly dysregulated canonical pathways by IPA (Figure 5F).

**Figure 5:**
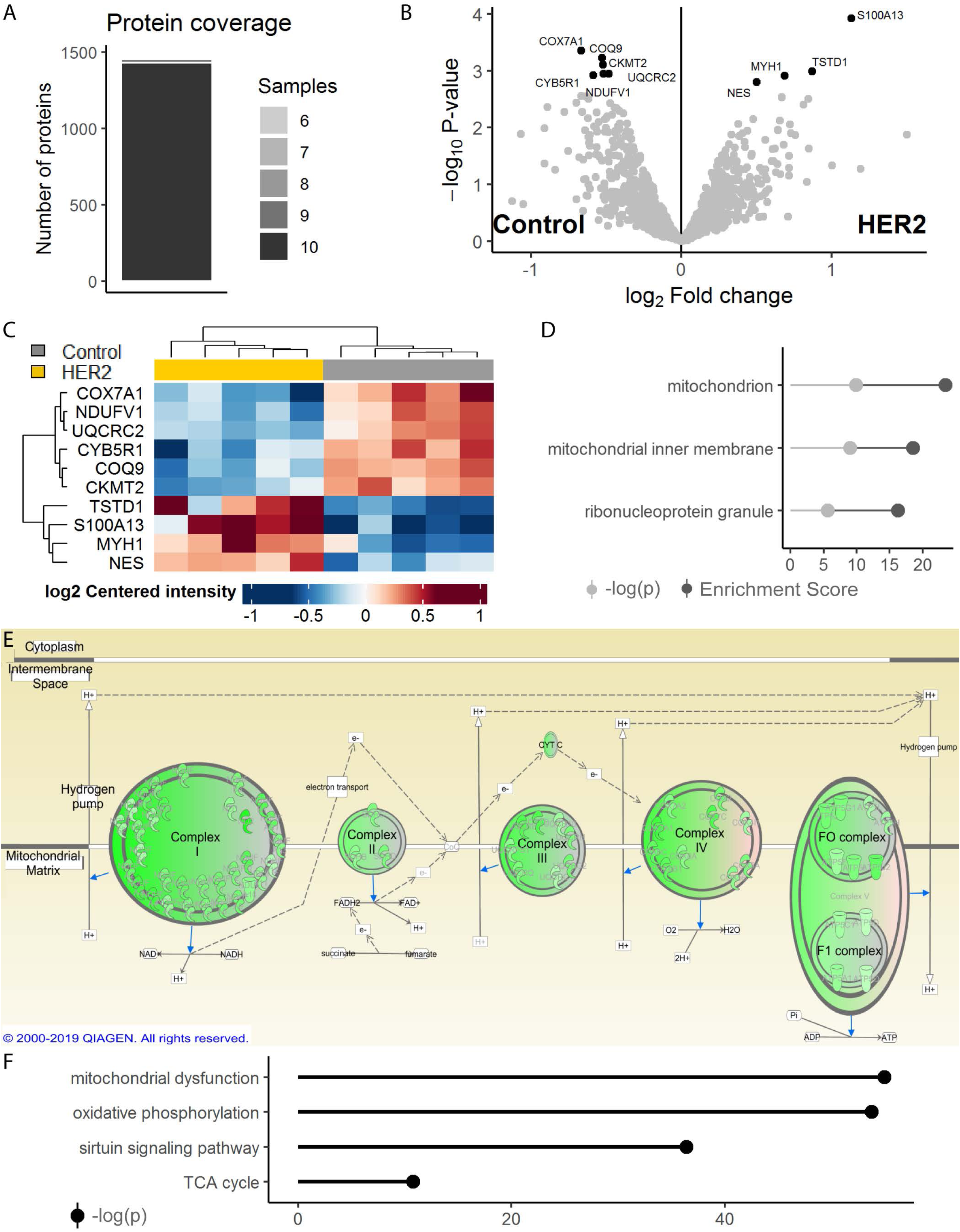
HER2 Patient Proteomics. Stacked bar chart representing the number of unique proteins detected by the number of samples (**A**, n=10 biopsies, n=1,555 unique proteins). Differential expression analysis of detected proteins, represented as a volcano plot with significantly differentially-expressed proteins identified in black (**B**). Heatmap of differentially-expressed proteins, with the ten columns representing the ten samples assayed (**C**). Enrichment analysis on the differentially-expressed proteins, querying Gene Ontology 2018 Cellular Components via Enrichr; top three results as ranked by Enrichment Score (**D**). IPA representation of the mitochondrial electron transport chain, with proteins detected at lower abundance notated in green (**E**). IPA-predicted pathway dysregulation using proteomic data (**F**).

### Subtype-independent DEG analysis and experimental validation

To identify potential mechanisms of muscle fatigue that are consistent across all molecular subtypes of BC, pathway-level enrichment analysis was conducted using a restricted dataset of genes that were significantly differentially expressed when comparing all BC patients to control (FDR < 0.10). In this analysis, we observed a strong signal indicating aberrant mitochondrial function across multiple databases queried. For example, three of the top four enriched pathways identified when querying the WikiPathways 2019 Human database included Electron Transport Chain System in Mitochondria (WP111), Oxidative Phosphorylation (WP623), and Mitochondrial Complex I Assembly (WP477, Figure 6A); and the three most enriched cellular components from the 2018 Gene Ontology project were all mitochondrial components (Figure 6B).

**Figure 6:**
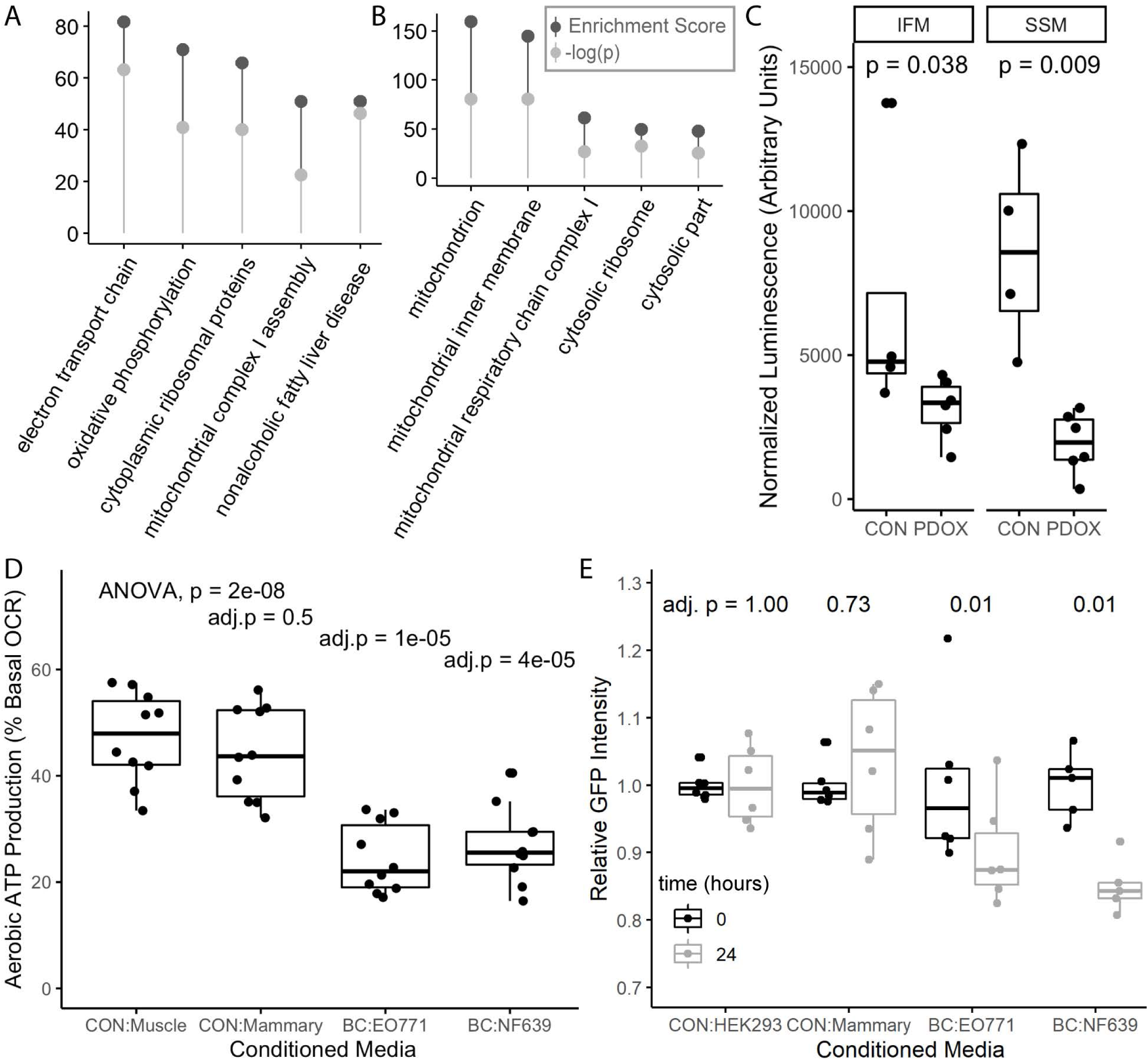
Subtype-Independent DEG Analysis and Experimental Validation. Enrichment analysis of DEGs, querying WikiPathways 2019 Human database: Electron Transport Chain System in Mitochondria (WP111), Oxidative Phosphorylation (WP623), and Mitochondrial Complex I Assembly (WP477). Top hits ranked by Enrichment Score (**A**). Enrichment analysis of DEGs, querying Gene Ontology 2018. Top hits ranked by Enrichment Score (**B**). Enrichment Score and p-value as provided by Enrichr. A –log(p) > 1.3 corresponds to p < 0.05 (**A, B**). ATP content in *quadriceps muscle* of control (n=4) and PDOX-bearing (n=6) mice as assessed by chemiluminescence in two mitochondrial subpopulations. ATP content in the PDOX mitochondria was compared to control using the Mann-Whitney U test (**C**). Aerobic ATP production as a percentage of basal oxygen consumption rate (OCR) in differentiated C2C12 myotubes treated for 48 hours with media conditioned by C2C12 (CON-Muscle), EpH4-EV (CON-Mammary), EO771 (BC), or NF639 (BC) cells (n=10 per group). Overall significance determined by one-way ANOVA followed by two-tailed Student’s t-tests with Bonferroni correction comparing each treatment group to CON-Muscle (**D**). Normalized GFP intensity in HEK293-PPRE-H2b-eGFP reporter cells at baseline and after treatment with conditioned media for 24 hours (n=6 for HEK293, EpH4-EV, and EO771; n=5 for NF639). Normalized GFP intensity values at 24 hours were compared to baseline measurements using paired samples two-tailed t-tests with Bonferroni correction of p-values (p. adj) (**E**).

To test this prediction of altered mitochondrial function in tissues distant from the primary tumor *in vivo*, we generated 6 female BC-PDOX mice and isolated live mitochondria from their skeletal muscle at euthanasia for quantification of ATP content. We found a significant reduction in ATP content within both of the two major skeletal muscle mitochondrial subpopulations, the interfibrillar mitochondria (IFM) and subsarcolemmal mitochondria (SSM), relative to female control animals (Figure 6C). *In vitro* assays were conducted to determine whether mitochondrial dysfunction in muscle is a direct response to BC-secreted factors or an indirect response mediated by other tissues. In support of a direct response, conditioned media from the luminal EO771 BC cell line and the HER2/neu overexpressing NF639 line significantly repressed aerobic ATP production in differentiated C2C12 myotubes, whereas media conditioned by either the normal mammary epithelial cell line EpH4-EV or C2C12 myoblasts did not (Figure 6D). To validate the role of the PPAR signaling proteins in mediating the systemic response to BC-secreted factors, conditioned media was isolated from EO771 and NF639 cells and applied to HEK293 cells stably expressing a PPAR-responsive promoter driving GFP expression. Media conditioned by both BC cell lines significantly repressed GFP intensity relative to reporter cell-conditioned media, while media conditioned by a normal mammary epithelial cell line did not alter GFP signal relative to control (Figure 6E). These data indicate that BC cells secrete a substance that is capable of directly influencing metabolic function in skeletal muscle, which may be related to a repression of PPAR-mediated transcriptional activity.

## DISCUSSION

BC-induced muscle dysfunction is a common problem of unclear etiology with few therapeutic options. Here we sought to identify possible mechanisms of fatigue that are generalizable across BC subtypes and are independent of treatment status by assessing statistical relationships between patients’ clinical characteristics and overall skeletal muscle gene expression.

In support of our laboratory’s previous publications ^14, 15^, we found that women with three molecular subtypes of BC, those being ERPR, TN, and TP BC, exhibit overall similarity in muscular gene expression. Remarkably, patients with tumors overexpressing HER2/neu in the absence of ER and PR expression exhibited a markedly different muscular gene expression profile. However, gene expression data from all subtypes pointed to similar pathway-level dysregulation, indicating that the mechanisms leading to muscle fatigue in BC patients may indeed be generalizable across subtypes. In assessing the commonalities across patients, we observed significant dysregulation of metabolic pathways in all groups of BC patients, and proteomic analysis of patients with HER2/neu-overexpressing tumors showed decreased abundance of nearly all proteins involved in the mitochondrial electron transport chain. Because mitochondrial density in skeletal muscle has been shown to correlate with abundance of ETC complex proteins and skeletal muscle oxidative capacity ^16^, it is likely that BC patients also have decreased mitochondrial density in their skeletal muscle as well as decreased oxidative capacity. We propose then that BC-secreted factors induce muscle dysfunction by abrogating oxidative capacity via alteration of mitochondrial biogenesis, mitophagy, or fission/fusion dynamics. Both *in vitro* and *in vivo* assays confirm that factors from BC cells alter skeletal muscle ATP content and/or aerobic ATP production, perhaps via dysregulation of the PPAR-signaling pathway.

Our laboratory’s interest in the PPAR family of proteins arose from our previous RNA-sequencing analysis of muscle from BC patients and PDOX-bearing mice, which our group found to recapitulate the clinical phenotype of increased muscle fatigue without muscle atrophy or bodyweight loss ^15^. These proteins have demonstrated roles in whole-body energy regulation, are critical regulators of mitochondrial function in multiple tissues, and are targets of multiple FDA-approved agents in the treatment of type 2 diabetes and hyperlipidemia. Among the three PPAR isoforms, we identified PPARG as a key regulator in BC-induced muscle dysfunction observed in PDOX mice ^15^. PPARG is a ligand-activated nuclear receptor that, upon activation by a variety of endogenous and synthetic lipids, forms complexes with retinoid X receptor (RXR) and cofactors such as the peroxisome proliferator-activated receptor-γ coactivator 1α (PGC1α), and stimulates transcription of downstream genes. This relationship with PGC1α is particularly relevant, as PGC1α is known to be a master regulator of mitochondrial biogenesis in several tissues, including skeletal muscle ^17^. PGC1α also participates in the regulation of other metabolic processes, including gluconeogenesis, muscle fiber-type specification, and control of antioxidant expression ^18–20^. Therefore, the interaction between PPARG and PGC1α clearly has the potential to impact muscle function through mitochondrial mechanisms. In addition to the potential development of muscle-intrinsic mitochondrial dysfunction, a systemic consequence of dysregulated PPARG is the development of insulin resistance (IR) ^21^, which is commonly associated with type 2 diabetes, obesity, and metabolic syndrome, and appears to have a bidirectional relationship with BC, with individuals with IR at greater risk of BC and BC survivors at an increased risk of IR ^22, 23^. We propose that the development of mitochondrial dysfunction and IR, secondary to muscular PPARG downregulation by BC, creates an environment that facilitates the development of muscle fatigue through several mechanisms, including decreased mitochondrial ATP production as well as dysregulated glucose and lipid metabolism. Pharmacological restoration of PPARG function results in the induction of a number of genes involved in insulin signaling, as well as glucose and lipid metabolism, and PPAR agonist drugs including the TZDs have shown a remarkable ability to restore insulin sensitivity in insulin-resistant conditions. Because our predictions have implicated many related metabolic pathways, we hypothesize that the development of BC induced muscle fatigue constitutes a pathology similar to type 2 diabetes/metabolic syndrome. If repression of PPAR signaling is indeed central to BC-induced muscle dysfunction, the numerous FDA-approved PPAR-agonists could address this unmet need in clinical oncology.

A particularly surprising result in the present study was that no clinical data aside from BC molecular subtype exhibited significant correlation with skeletal muscle gene expression, including TNM staging and history of chemotherapy, radiation, or immunotherapy. Additionally, patients in the HER2 group consistently exhibited a decreased abundance of mitochondrial proteins in their muscle tissue, despite significant differences in prior treatments, and importantly, we observed these responses consistently across all five biopsies from this patient group, despite significant differences in anti-cancer treatment history. At the date of biopsy collection, one patient was entirely treatment naïve, one patient had completed chemotherapy for BC 10 years prior, and the remaining 3 patients received different multi-agent neoadjuvant chemotherapy in the months prior to surgery. Therefore, we propose that changes in skeletal muscle physiology seen in BC are due to tumor-derived factors rather than side-effects of therapies or other patient-specific factors. This hypothesis is supported by our *in vitro* conditioned media experiments where tumor-derived factors directly repressed mitochondrial respiratory capacity in differentiated muscle cells. Additionally, neither weight loss nor body composition were predictive of skeletal muscle gene expression, suggesting that gene expression changes observed in BC patients are unrelated to muscle atrophy or cachexia and may instead be reflective of muscle dysfunction.

Because albumin is often used by physicians as a measure of patients’ nutritional status and mortality risk ^24–27^, we assessed serum albumin in our patient cohort and assessed its relationship to skeletal muscle gene expression and weight change. In our analysis, serum albumin was not found to be predictive of weight change, and in a larger sample was found to be only weakly predictive of cachexia risk. Yet, it was unexpectedly predictive of skeletal muscle gene expression. This indicates that serum albumin may be a useful biomarker of muscle function in the absence of cachexia, a possibility supported by previous literature connecting serum albumin with muscle strength ^28^ and insulin resistance ^29^ in other clinical contexts. Prospective studies directly addressing the relationship between skeletal muscle function, gene expression, and serum albumin would be required to validate the clinical utility of serum albumin in identifying those at risk of cancer-induced muscle dysfunction.

In line with our previous publications ^14, 15^, we report that transcriptional responses to the ERPR, TP and TN subtypes of BC are similar in terms of skeletal muscle gene expression, while muscle biopsies from patients with tumors overexpressing HER2/neu in the absence of ER and PR exhibit an unusual degree of uniqueness in terms of gene expression. We identified a strong signal for BC-induced mitochondrial dysfunction in BC patients, PDOX-bearing animals, and in *in vitro* assays, and proteomic analysis showed decreased protein abundance of nearly all components of the mitochondrial electron transport chain in muscle biopsies from patients with HER2/neu-overexpressing tumors relative to control. Further we found no relationship between various BC-related treatments (surgery, chemotherapy, radiotherapy) and changes in skeletal muscle gene expression. However, serum albumin was predictive of skeletal muscle gene expression without being predictive of weight loss, suggesting that serum albumin may be a useful indicator of BC-induced skeletal muscle dysfunction. Overall, these data indicate that all BC subtypes induce dysfunction in mitochondrial respiration in skeletal muscle independent of molecular subtype, and this effect appears to be independent of anti-cancer treatments. These findings call for prospective studies assessing interventions to support skeletal muscle function in the presence of BC-induced mitochondrial dysfunction.

## METHODS

### Patient information

Conduct of research involving human patients at West Virginia University is guided by the principles set forth in the Ethical Principles and Guidelines for the Protection of Human Subjects of Research (Belmont Report) and is performed in accordance with the Department of Health and Human Services policy and regulations. A total of 71 female surgical patients provided informed consent for inclusion in this study (control n=20; BC n=51). Information regarding patient selection and consenting process have been previously published ^15^. Women with BC provided muscle biopsies from the *pectoralis major muscle* intraoperatively at the time of mastectomy, and control patients provided *pectoralis major muscle* samples intraoperatively during other breast surgeries. Women with BC were classified into four molecular subtypes based on immunohistochemical staining of their primary tumors: ERPR (n=20), HER2 (n=9), TN (n=11), or TP (n=11).

Information on BMI at multiple time points was collected in 12 control and 50 BC patients. The mean number of days a consented patient was followed was 170 +/- 214 (control median=170, ERPR median=36, HER2 median=146, TP median=140, TN median=180). Detailed body composition analyses were acquired in 8 control and 35 BC patients using a bioelectrical impedance scale (Tanita: model SC-240). Serum albumin levels were acquired in 3 control and 49 BC patients (ERPR n=18; HER2 n=9; TP n=11; TN n=10). For RNA-Seq analyses, biopsies were used from 10 control and 33 BC patients (ERPR n=10; HER2 n=5; TP n=9; TN n=9).

In a separate analysis of de-identified electronic medical records (EMRs) from female BC patients, body mass and standing height data were acquired for 5,201 individuals for calculation of BMI. Within this population, final analyses were completed in 3,001 patients having at least two measurements for weight in addition to at least one record for serum albumin.

### RNA-Sequencing

*Pectoralis major muscle* biopsies were acquired intraoperatively and stored in Invitrogen RNAlater Solution (Thermo Fisher, San Jose, CA) overnight at 4°C and then at −80°C until processing. RNA was isolated, assessed for quality, and utilized to construct libraries for RNA-Seq as previously reported ^15^. Completed libraries were sequenced on one lane of the HiSeq 1500 with PE50 bp reads. Subsequently, Salmon was used for transcript-level abundance estimation, with both gcBias and seqBias set, and libType A ^30^. Transcript-level abundance estimates were summarized to the gene-level using *tximport* ^31^. The resulting counts matrix was scaled to library size using *edgeR* ^32^ and filtered to remove genes without a counts-per-million (CPM) value > 1 in 3 or more samples. Log-transformed CPM values for the 10,000 genes with highest variance were used as input for heatmap creation using *gplots* ^33^. CPM values were log-transformed and scaled prior to distance matrix computation. The resulting distance matrix was then input for classical multidimensional scaling with k=3 and visualized in 3-dimensions using *car* ^34^. Differential gene expression analysis was conducted using *DESeq2* ^35^.

### Proteomics. Sample Preparation

*Pectoralis major muscle* biopsies from n=5 female patients with HER2/neu-overexpressing BC and n=5 control breast surgical patients were acquired intraoperatively and stored in RNA-later at −80°C until processing. Tissue fragments weighing approximately 50mg were sent on dry ice to the Mass Spectrometry and Proteomics Resource Laboratory at Harvard University and processed according to established protocols ^36, 37^. Briefly, biopsies were lysed in Covaris^®^ microTUBE-15 (Woburn, MA) microtubes with Covaris^®^ TPP buffer, using the Covaris S220 Focused-ultrasonicator instrument with 125W power over 180s with 10% max peak power. Samples were then chloroform/methanol precipitated, filtered, reduced, alkylated, and finally digested overnight at 38°C in a solution containing triethylammonium bicarbonate and Promega^®^ Sequencing Grade Trypsin. Digested peptides were then incubated with tandem mass tag, using different tags for each sample.

### LC-MS/MS

The 10 samples were pooled in equal amounts and fractionated into 10 fractions. LC-MS/MS was performed on an Orbitrap Lumos (Thermo Fisher) equipped with EASYLC1000 (Thermo Fisher). Peptides were separated onto a 100 µm inner diameter microcapillary column packed first with C18 Reprosil resin (5 µm, 100 Å, Dr. Maisch GmbH, Germany) followed by analytical column of Reprosil resin (1.8 µm, 200 Å, Dr. Maisch GmbH, Germany). Separation was achieved through applying a gradient from 5– 27% acetonitrile in 0.1% formic acid over 90 min at 200 nl**·**min^−1^. Electrospray ionization was enabled through applying a voltage of 1.8 kV using a homemade electrode junction at the end of the microcapillary column and sprayed from fused silica pico tips (New Objective, MA). The LTQ Orbitrap Lumos was operated in data-dependent mode for the mass spectrometry methods. The mass spectrometry survey scan was performed in the Orbitrap in the range of 395 –1,800 m/z at a resolution of 6 × 10^4^, followed by the selection of the twenty most intense ions (TOP20) for collision induced dissociation (CID) in the Ion trap using a precursor isolation width window of 2 m/z, AGC setting of 10,000, and a maximum ion accumulation of 200 ms. Singly charged ion species were not subjected to CID fragmentation. Normalized collision energy was set to 35 V and an activation time of 10 ms. Ions in a 10 ppm m/z window around ions selected for MS2 were excluded from further selection for fragmentation for 60 s. The same TOP20 ions were subjected to higher-energy collisional dissociation (HCD) MS2 event in Orbitrap part of the instrument. The fragment ion isolation width was set to 0.7 m/z, AGC was set to 50,000, the maximum ion time was 200 ms, normalized collision energy was set to 27V and an activation time of 1 ms for each HCD MS2 scan.

### Mass spectrometry analysis

Raw data were submitted for analysis in Proteome Discoverer 2.2 (Thermo Scientific). Assignment of MS/MS spectra was performed using the Sequest HT algorithm by searching the data against a protein sequence database including all entries from Uniport_Human2016_SPonly database as well as other known contaminants such as human keratins and common lab contaminants. Sequest HT searches were performed using a 20 ppm precursor ion tolerance and requiring each peptides N-/C termini to adhere with trypsin protease specificity, while allowing up to two missed cleavages. 6-plex TMT tags on peptide N termini and lysine residues (+229.162932 Da) was set as static modifications while methionine oxidation (+15.99492 Da) was set as variable modification. A MS2 spectra assignment false discovery rate (FDR) of 1% on both protein and peptide level was achieved by applying the target-decoy database search. Filtering was performed using a Percolator ^38^ (64bit version). For quantification, a 0.02 m/z window centered on the theoretical m/z value of each the six reporter ions and the intensity of the signal closest to the theoretical m/z value was recorded. Reporter ion intensities were exported using Proteome Discoverer 2.2.

### BC-PDOX skeletal muscle mitochondria metabolic analysis

Animal experiments were approved by the WVU Institutional Animal Care and Use Committee. BC-PDOX mice were created by implanting human BC tumor fragments into the mammary fat pad of female NOD.CG-Prkdscid Il2rgtm1 Wjl/SzJ/ 0557 (NSG) mice (n=6), as described previously ^15^. PDOX-bearing animals were euthanized approximately 30 days after reaching a tumor volume of 200 mm^3^. Control female NSG mice of similar age (n=4) were euthanized at the same time as tumor-bearing animals and tissues were processed identically in both groups. Immediately after death, both quadriceps muscles from each mouse were quickly removed and interfibrillar (IFM) and subsarcolemmal (SSM) mitochondria were isolated separately according to previously described methods ^39–42^, combining the two muscles to obtain sufficient tissue for downstream applications. Mitochondrial isolates were stored at −80°C until analysis. ATP content was quantified in each mitochondrial subpopulation using the ENLITEN® ATP Assay System Bioluminescence Detection Kit (Promega, Wisconsin, USA) according to manufacturer’s protocol with minor modifications, as follows. Mitochondrial isolates were lysed in 1% trichloroacetic acid for 5 minutes then diluted 1:10 in 0.1 mol**·**L^-1^ tris base (pH = 7.8). The resulting samples were loaded into a black-walled microplate, in duplicate, and mixed 1:1 with luciferase/luciferin reagent. Luminescence intensity was immediately read using the FlexStation® 3 Multi-Mode Microplate Reader (Molecular Devices®, California, USA). Luminescence intensity values were blank-corrected and normalized to account for differing protein concentrations, which were quantified using the DC™ Protein Assay (Bio-Rad, California, USA).

### Cell culture

All cell lines were obtained from ATCC (Virginia, USA) and cultured in Gibco DMEM (Thermo Fisher, Massachusetts, USA) supplemented with 10% heat inactivated fetal bovine serum (Atlanta Biologicals, Georgia, USA) and Gibco penicillin/streptomycin (Thermo Fisher) at 37°C with 6% CO_2_. Cell lines utilized include EpH4-EV (immortalized normal murine mammary epithelium), EO771 (murine luminal BC), NF639 (murine HER2/neu-overexpressing BC), HEK293 (human embryonic kidney), and C2C12 (murine myoblasts).

### *In vitro* conditioned media (CM) metabolic analysis

C2C12 cells were plated into Agilent Seahorse XF24 (Agilent Technologies, California, USA) plates and differentiated by confluence for 3 days. Meanwhile, EpH4-EV, EO771, NF639, and C2C12 cells were plated at approximately 15% confluence in separate 10cm dishes for 72 hours. The 72-hour CM was then removed from all cell lines, centrifuged at 1,500 RPM for 10 minutes, and then the supernatants were collected, diluted 1:3 in fresh growth media, and applied to the differentiated C2C12 cells in Seahorse assay plates for 48 hours (n=10 wells per treatment condition) prior to conducting the Agilent Seahorse XF Cell Mito Stress Test protocol according to manufacturer’s instructions.

### *In vitro* CM PPAR-reporter assays

HEK293 cells were transfected with PPRE-H2b-eGFP ^43^ (Addgene #84393) using Invitrogen Lipofectamine 3000 (Thermo Fisher), selected with 500 ng**·**uL^-1^ Gibco geneticin (Thermo Fisher) for 20 days, and flow-sorted to select the cells expressing GFP. The resulting HEK293-PPRE-H2b-eGFP cell line was plated at approximately 15% confluence in a 24-well plate. The following day, cells were imaged using the BioTek Cytation 5 Cell Imaging Multi-Mode Reader (Agilent Technologies) to collect baseline GFP intensity, and 72-hour conditioned media from HEK293, EpH4-EV, EO771, and NF639 well lines was applied to the 24-well plate, using individual wells as biological replicates (n=6 wells per treatment condition). Cells were then incubated in the conditioned media under normal culture conditions for 24 hours, at which point GFP intensity was measured using identical imaging settings as the baseline collection. Mean cellular GFP intensity per well was calculated using Gen5 Microplate Reader and Imager Software (Agilent Technologies) after background flattening and thresholding, which were set consistently across all images.

### Statistical Analyses. Clinical information

Patients’ trends of weight change over time were calculated for each individual patient by fitting a simple linear regression line to their weight at each date in the EMR, normalized such that each patient’s first weight record equaled 100, resulting in a slope representing their approximate percentage of body weight change per day. 19 patients were identified as outliers in terms of daily weight change. 13 of these patients were excluded due to having a limited observation period (< 15 days). Each patient’s first albumin measurement was then obtained and the rate of daily weight change was regressed on the patients’ first albumin measurements. Pearson’s correlation coefficients and p-values were calculated and plotted using *ggpubr* ^44^ in R v3.6.1 ^45^.

Logistic regression analysis was utilized to determine whether serum albumin was predictive for a rate of weight change consistent with cachexia (i.e. < −0.027% per day to reach 5% weight loss in 6 months ^1, 2^). Omnibus model fit was assessed by Chi-square test and effect size was calculated using Nagelkerke’s pseudo-R^2^. The receiver operating characteristic curve (ROC) test was conducted, with preferred sensitivity and specificity > 0.7. No point on the ROC satisfied these conditions. The fitted logistic regression model was used to predict whether each patient would exhibit a rate of weight loss consistent with cachexia, and these predictions were compared to the actual data to create a confusion matrix for determination of sensitivity, specificity, and positive predictive value.

### RNA-Seq and clinical correlates

Clinical data were assessed against the 3-dimensional distance matrix representing overall skeletal muscle gene expression in the context of a multivariate regression model. Each clinical variable available was regressed on the three outcome variables and model fit was assessed via MANOVA and Pillai’s trace. Forward selection was then applied to combine the variables with smallest p-values for Pillai’s trace into a final model, with final selection considerations including statistical significance for MANCOVA using Pillai’s trace and Wilks’ lambda, maximizing Pillai’s trace test statistic, and minimizing Wilks’ lambda test statistic.

Differential gene expression analysis was conducted using *DESeq2* ^35^. Input data consisted of transcript-level abundance estimates from Salmon summarized to the gene level using *tximport* ^31^. Two differential expression analyses were run: one comparing the group of BC patients to control patients, and another comparing the BC patients by subtype to control patients, with the null hypothesis rejected at FDR < 0.10. Because molecular subtype was the only variable that yielded statistical significance for predicting gene expression in the multivariate regression model described above, no clinical characteristics were assessed in the differential expression analysis.

### Proteomics

Differential protein expression analysis was conducted using *DEP* ^46^. Protein expression values were first filtered to remove known contaminants and proteins with missing expression values in more than 1 sample per group, then normalized, and then background-corrected using variance stabilizing transformation. Remaining missing values were imputed using k-nearest neighbor, after determining that the small number of missing values were likely missing at random. Differential expression analysis using linear models and empirical Bayes statistics was then conducted on the imputed dataset, with the null hypothesis rejected at FDR < 0.05.

### Enrichment analyses

Input data for Qiagen’s Ingenuity Pathway Analysis (IPA) consisted of differential expression values and false discovery rate-adjusted p-values (FDR) from proteomic and RNA-sequencing data (FDR < 0.5). Within IPA, all human data sources were queried, with confidence threshold set to include only experimentally observed effects and those predicted with high confidence. Input data for Enrichr ^47, 48^ analyses included only significantly differentially expressed genes and proteins. Data presented in this article reflect analyses conducted in Enrichr and IPA between May 15^th^ and June 15^th^ of 2019.

### In vitro and in vivo validation assays

ATP content in each mitochondrial subpopulation in the PDOX muscle were compared to control using the Mann-Whitney U test. The null hypothesis was rejected at p < 0.05. In the conditioned media metabolic experiments, one-way ANOVA was used to compare the rate of oxygen consumed in ATP production as a percentage of basal oxygen consumption between the four conditioned media treatment groups followed by two-tailed Student’s t-tests with Bonferroni correction comparing each treatment group to CON-Muscle. The null hypothesis was rejected at p < 0.05. This experiment was conducted as reported twice with similar results, and results from the first analysis are reported. In the PPAR-responsive reporter assays, mean GFP intensities in each well were normalized to account for differences in baseline GFP expression between wells. Normalized GFP intensities at 24 hours were compared to normalized baseline measurements using a paired samples two-tailed t-test by treatment group with Bonferroni correction for multiple comparisons. The null hypothesis was rejected at p < 0.05. This experiment was conducted as reported twice with similar results, with results from the first analysis reported. This analysis includes n=5 for NF639-treated cells due to a technical problem during baseline image capture that resulted in the loss of one image.

### Chi-Square analyses

First, the actual number of unique and shared DEGs and dysregulated pathways were counted for each subtype. For example:

HER2-unique = genes differentially-expressed only in HER2 patients

HER2-2-Group = HER2 ∩ ERPR + HER2 ∩ TP + HER2 ∩ TN;

HER2-3-Group = HER2 ∩ ERPR ∩ TP + HER2 ∩ ERPR ∩ TN + HER2 ∩ TN ∩ TP.

Then, expected numbers of DEGs and dysregulated pathways were calculated under the null hypothesis that DEGs and dysregulated pathways are independent of BC subtype. Chi-square statistics were calculated using the difference between observed and expected numbers of DEGs and dysregulated pathways. On 6 degrees of freedom, the Chi-square critical value is 12.592 for statistical significance at α = 0.05. Bar plots represent the log-fold change in number of DEGs or dysregulated pathways compared to expected values, by category, and actual Chi-square statistics for each category are also provided.

### Box-and-whisker plots

The width of the box represents the interquartile range (IQR) and whiskers extend to the single most extreme measurement in both directions, unless the most extreme measurement is considered an outlier, in which case the most extreme value is represented by two dots. Median values are represented by the horizontal line through each boxplot. Outliers in this context are defined as values more extreme than 1.5 x IQR.

## Acknowledgements

This research was supported by the following: National Institute of General Medical Sciences of the National Institutes of Health under Award Number P20GM121322 (Lockman), American Cancer Society Institutional Research Grant 09-061-04 (Pistilli), the WVCTSI U54GM104942 (Hodder). Authors would like to acknowledge the following WVU Core Facilities for contributing to this work: Genomics Core Facility; Flow Cytometry and Single Cell Core Facility (S10OD016165); Preclinical Tumor Models Core Facility (CA148671, Pugacheva); Mitochondria Core of the WVU Stroke CoBRE (P20GM109098); Mitochondria, Metabolism and Bioenergetics group (R01 HL-128485; Hollander and the Community Foundation for the Ohio Valley Whipkey Trust). Additional acknowledgements to the Mass Spectrometry and Proteomics Resource Laboratory at Harvard University for proteomics analyses and to Dr. Metheny-Barlow at Wake Forest University for providing EO771 cells.

---

**Supplemental Figure 1.** Scaled, log-transformed protein expression correlated with scaled, log-transformed RNA expression in a gene-wise fashion, presented for 8/8 patients with both RNA-seq and proteomic analyses.

