## Supplementary figures and images for "Skeletal muscle reprogramming by breast cancer regardless of treatment history or tumor molecular subtype"

### Supplemental Figure 1

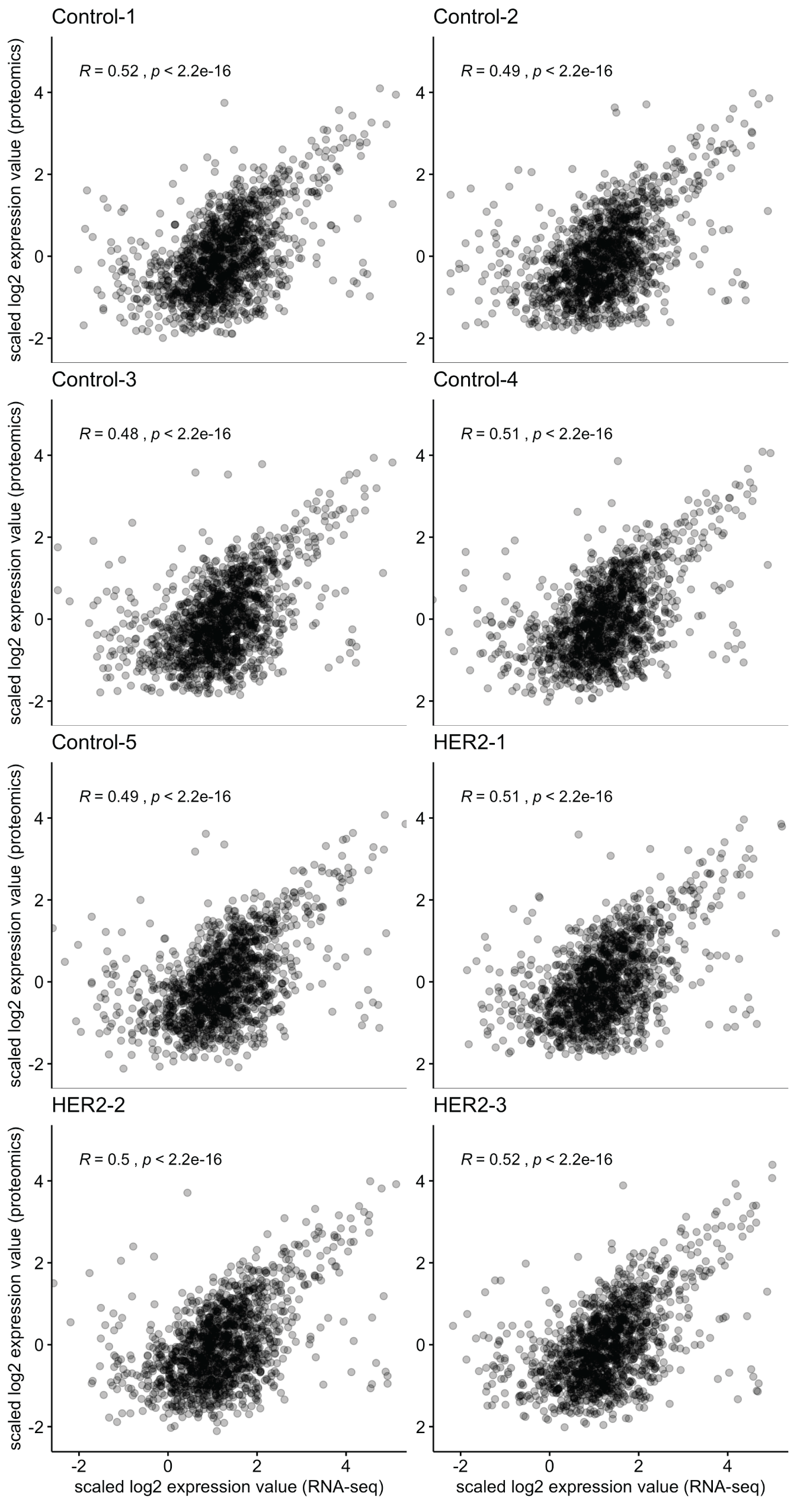
